# Interneuron loss and microglia activation by transcriptome analyses in the basal ganglia of Tourette syndrome

**DOI:** 10.1101/2024.02.28.582504

**Authors:** Yifan Wang, Liana Fasching, Feinan Wu, Anita Huttner, Sabina Berretta, Rosalinda Roberts, James F. Leckman, Alexej Abyzov, Flora M. Vaccarino

**Affiliations:** Department of Quantitative Health Sciences, Center for Individualized Medicine, Mayo Clinic, Rochester, MN 55905, USA; Child Study Center, Yale University, New Haven, CT 06520, USA; Department of Pathology, Yale University, New Haven, CT 06520, USA; McLean Hospital, Harvard Medical School, Belmont, MA, 02478, USA; Department of Psychiatry and Behavioral Neurobiology, University of Alabama at Birmingham, Birmingham, AL 35294, USA; Department of Neuroscience, Yale University, New Haven, CT 06520, USA; Yale Kavli Institute for Neuroscience, New Haven, CT 06520, USA

## Abstract

Tourette syndrome (TS) is a disorder of high-order integration of sensory, motor, and cognitive functions afflicting as many as 1 in 150 children and characterized by motor hyperactivity and tics. Despite high familial recurrence rates, a few risk genes and no biomarkers have emerged as causative or predisposing factors. The syndrome is believed to originate in basal ganglia, where patterns of motor programs are encoded. Postmortem immunocytochemical analyses of brains with severe TS revealed decreases in cholinergic, fast-spiking parvalbumin, and somatostatin interneurons within the striatum (caudate and putamen nuclei). Here, we performed single cell transcriptomic and chromatin accessibility analyses of the caudate nucleus from 6 adult TS and 6 control post-mortem brains. The data reproduced the known cellular composition of the adult human striatum, including a majority of medium spiny neurons (MSN) and small populations of GABAergic and cholinergic interneurons. Comparative analysis revealed that interneurons were decreased by roughly 50% in TS brains, while no difference was observed for other cell types. Differential gene expression analysis suggested that mitochondrial function, and specifically oxidative metabolism, in MSN and synaptic function in interneurons are both impaired in TS subjects. Furthermore, such an impairment was coupled with activation of immune response pathways in microglia. Also, our data explicitly link gene expression changes to changes in cis-regulatory activity in the corresponding cell types, suggesting de-regulation as a factor for the etiology of TS. These findings expand on previous research and suggest that impaired modulation of striatal function by interneurons may be the origin of TS symptoms.

## Introduction

Tourette syndrome (TS) is a common disorder afflicting as many as 1 in 150 children (1). The main symptoms are motor and vocal tics, such as eye-blinking or throat clearing, that are performed out-of-context and in a repetitive fashion (2). TS has one of the highest familial recurrence rates among complex neuropsychiatric disorders(3); yet few risk genes meeting genome-wide statistical thresholds and no biomarkers have emerged as causative or predisposing factors, despite the efforts of international consortia and the combined examination of nearly 5,000 TS and 10,000 normal control (NC) blood samples (3).

TS is a disorder of high-order integration of sensory, motor, and cognitive functions. Patterns of motor programs are encoded in the basal ganglia (BG), a set of interconnected nuclei situated deep within the cerebral hemispheres (**Fig. 1A**). The BG are activated during tics in TS (4, 5). Neuroimaging data suggest that the caudate and putamen (referred to as striatum) are smaller in TS (6, 7). Notably, caudate volumes in childhood are predictive of TS severity in adulthood (8) and are reduced in the most severely affected twin in monozygotic twin discordant for TS (9). Postmortem immunocytochemical analyses in the striatum of adult individuals with severe TS revealed decreases in cholinergic interneurons (CH-IN) and in two GABAergic interneurons subtypes: parvalbumin interneurons (PV-IN) and somatostatin/nitric oxide synthase/neuropeptide Y interneurons (SST/NOS/NPY-IN), suggesting decreased generation, loss or hypofunction of these cell types (10-12).

**Figure 1.**
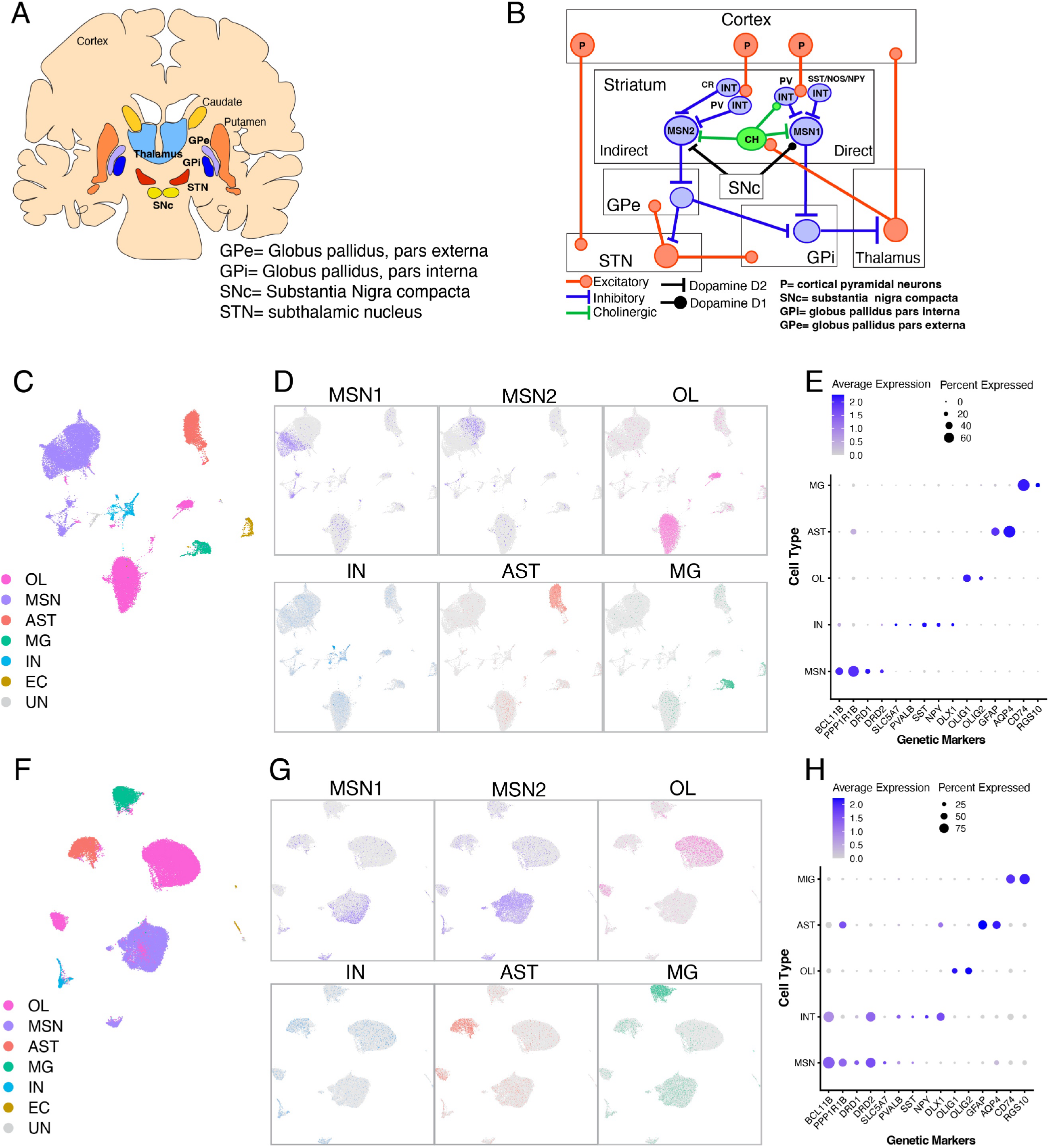
Clustering cells based on snRNA-seq and snATAC-seq data and defining cell types. **A)** Outline of basal ganglia (BG) anatomy in a coronal human brain section. **B)** Wiring diagram of BG circuitry. **C)** UMAP clustering of nuclei from 12 brains based on snRNA-seq data. Clusters (see Supplementary Fig. 1) are grouped into six cell types. **D)** UMAP plots colored by marker expression for the six cell types. Markers in order: DRD1 (for MSN1), DRD2 (for MSN2), OLIG1 (for OL), a combination of IN markers, AQP4 (for AST), CD74 (for MG). **E)** snRNA marker expression in cell types. Color of dots: average expression; size: percent of nuclei expressing the gene. **F)** UMAP clustering of nuclei from 12 brains based on snATAC-seq data. Clusters are annotated based on label transfer from snRNA-seq clusters. **G)** UMAPs from ATAC-seq data colored by marker expression for different cell types. Markers in order: DRD1 (for MSN1), DRD2 (for MSN2), OLIG1 (for OL), DLX1 (for IN), GFAP (for AST), RGS10 (for MG). **H)** snATAC marker expression in cell types. Color of dots: average expression; size: percent of nuclei expressing the gene. For details for all genetic markers see **Supplementary Table 6**.

The BG tonically inhibit the ventral and intralaminar thalamic nuclei through their common output, the globus pallidus pars interna/substantia nigra reticulata (GPi/SNr). The striatum receives converging inputs from the cortex, thalamus and midbrain dopamine neurons (**Fig. 1B**). The striatal medium spiny neurons (MSNs) form two major pathways, the indirect and direct pathway, which have opposite behavioral effects (13). The direct pathway inhibits the GPi/SNr, resulting in disinhibition of thalamic neurons and facilitation of motor behavior; conversely, the indirect pathway excites the GPi/SNr, inhibiting motor behavior. Striatal interneurons exert important modulatory roles on these two pathways. The fast-spiking, PV-IN respond with short latency to cortical stimulation (14-16) and exert feed-forward inhibition preferentially on MSNs of the direct pathway (17, 18). The CH-IN receive input from the intralaminar nuclei of the thalamus (19) and modulate the activity of IN and MSN (20) (**Fig. 1B**). Hence, the reported decreases in interneuron number in TS(10, 11) should hinder the inhibitory influence of thalamus and cortex upon the activity of the striatal direct pathway. This notion is supported by work in animal models, as selective ablation of CH-IN, or combined ablation of PV-IN and CH-IN in mouse, causes repetitive stereotypic behavior and increases functional connectivity between frontal cortical regions and the motor region of the striatum (21-23). Since previous work on striatal interneurons in TS was carried out primarily by immunocytochemical analyses, here we sought to advance our understanding of the pathophysiology of TS by performing single cell transcriptomic and chromatin accessibility analyses to both confirm the previous findings on the RNA level and at the same time perform a comprehensive characterization of the cellular composition and gene expression of the BG in TS individuals.

## Results

### Identification and annotation of major cell types

Human brain basal ganglia frozen brain samples from the head of the caudate nucleus were obtained from 12 individuals (6 patients with Tourette syndrome, 6 controls, paired by age and sex; **Supplementary Table 1**) and nuclear fractions were processed for single cell transcriptome analyses. For 3 pairs, we used stand-alone single-nuclei RNA-seq (snRNA-seq) and single nuclei ATAC-seq (snATAC-seq), while the remaining 3 pairs were processed using the multiome snRNA-seq and snATAC-seq platform from 10x, which allows deriving transcriptomic and epigenomic data from the same cell. We uniformly processed all 12 samples and concatenated the data from both techniques for downstream analysis (**Methods**).

After data QC, we retained 43,713 high-quality single nuclei whose expression profiles were clustered by similarity into 26 clusters and visualized via UMAP (**Supplementary Figure 1A**) with roughly the same distribution of clusters across the 12 brains (**Supplementary Figure 1B**). Clusters were grouped into 6 major cell types using known markers and public datasets(24, 25) (**Figure 1C**,**D**,**E**; **Methods**). We annotated 18,498 (42%) medium spiny neurons (MSN), 1,492 (3%) interneurons (IN), 5,394 (12%) astrocytes (AST), 15,563 (36%) oligodendrocytes (OL), 1,496 (3%) microglia (MG), 719 (2%) endothelial cells (EC), while 551 cells (1%) remained unidentified (UN) (**Supplementary Table 2**,**3, Supplementary Figure 2**). Overall, this cell distribution roughly corresponded to known proportions of cell types in human striatum, e.g., neurons account for ∼45% of all cells, while MSN account for ∼93% of all neurons, and IN account for ∼7% of all neurons. The MSN were composed of two subclusters, those expressing dopamine receptor 1 (DRD1)-MSN1 (55%) and those expressing dopamine receptor 2 (DRD2)-MSN2 (45%) (**Supplementary Figure 3**) which represent projection neurons of the direct and indirect pathways, respectively, distinguishable by specific gene expression profiles and each projecting *in vivo* to a separate set of targets, the globus pallidus pars interna and globus pallidus pars externa (**Fig. 1B**) (26).

**Figure 2.**
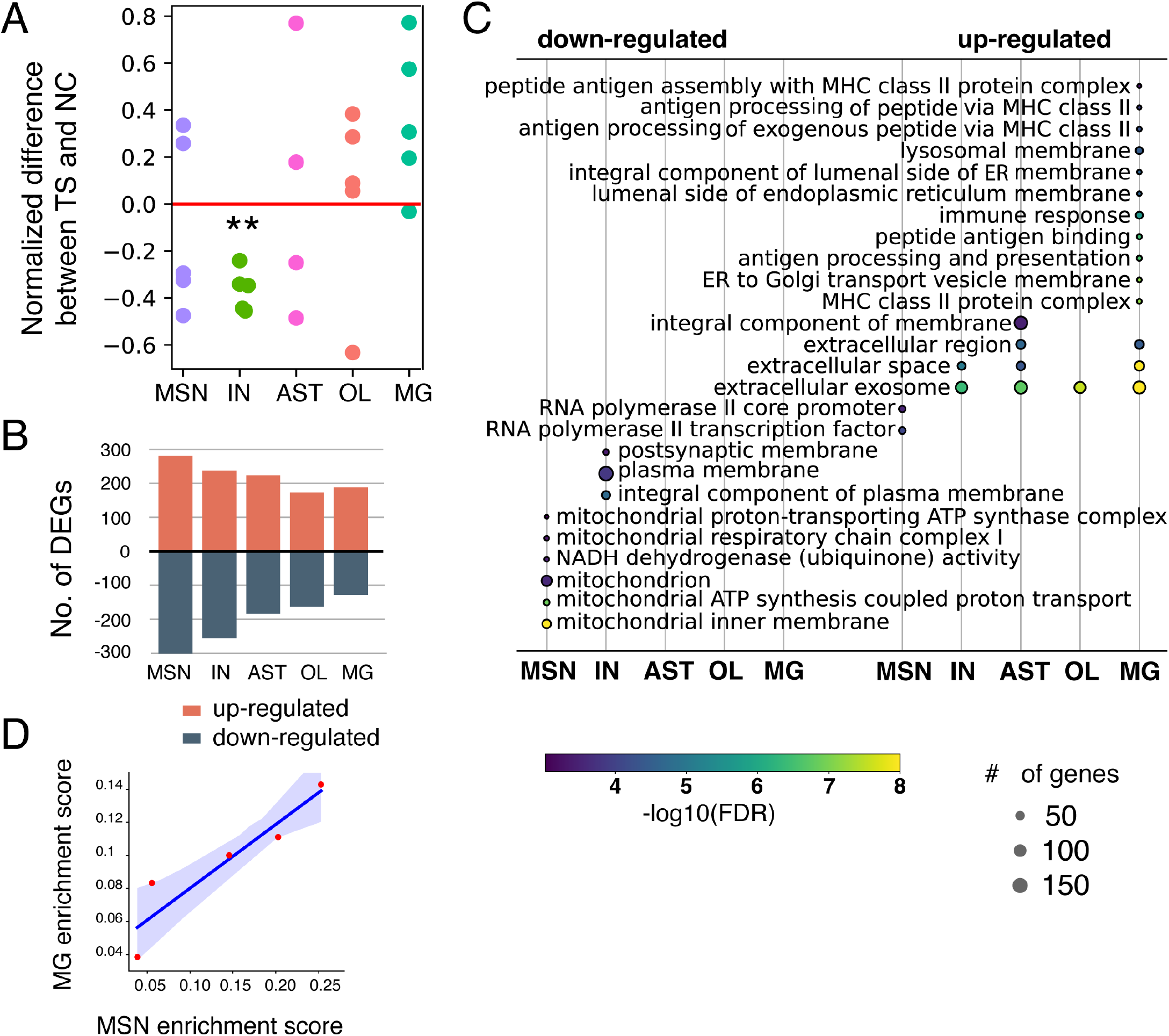
Transcriptomic profiles of Tourette syndrome brains compared to normal controls. **A)** Normalized difference in cell type proportions comparing TS to NC brains assessed using snRNA-seq data. **p-value < 0.001 by paired t-test. **B)** Number of differentially expressed genes in each cell types comparing TS to NC brains. **C)** GO terms enrichment for differentially expressed genes (up-regulated and down-regulated) in TS vs NC brains. y-axis: enriched GO terms; x-axis: cell types. The size of a circle indicates the number of genes in the enriched GO term, while color corresponds to -log10(FDR) for the enrichment. GO terms were ordered based on the p-value in each cell type. **D)** Correlation across 5 pairs of brains (red dots) between enrichment in mitochondrial function related pathways in MSN (x-axis) and immune response-related pathways in MG (y-axis).

**Figure 3.**
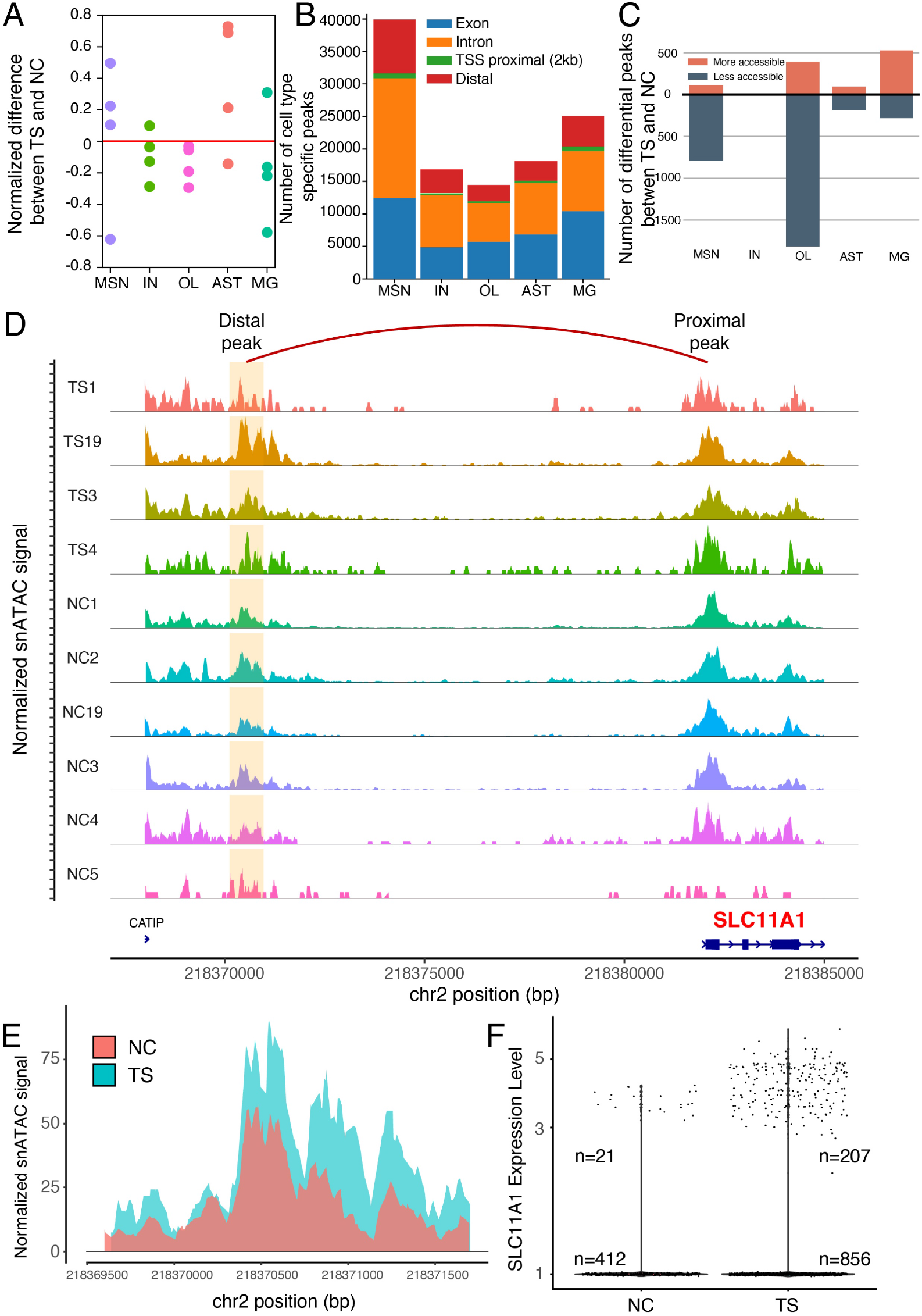
Chromatin accessibility profiles of Tourette syndrome brains compared to normal controls. **A)** Difference in cell type proportions comparing TS to NC brains assessed using snATAC-seq data. Y-axis: normalized percentage difference between TS and NC; x-axis: cell types. **B)** Number of snATAC-seq peaks across genome annotations. Y-axis: number of peaks; x-axis: cell types. **C)** Number of differentially accessible peaks (DAP) in TS as compared to NC brains. Y-axis: number of peaks; x-axis: cell types. **D)** Up-regulation of the SLC11A1 transcript in MG correlates positively with a more accessible distal DAP in TS brains. x-axis: genome coordinates; y-axis: aggregated normalized snATAC-seq signal in MGs of each brain; orange areas: snATAC-seq distal peaks co-accessible with snATAC-seq peaks proximal to the SLC11A1 promoter; red arches: predicted co-accessible peak pairs. **E)** More accessible snATAC-seq distal peaks combined in all TS vs all NC brains which are co-accessible with SLC11A1 promoter (adjusted p-value < 0.001, log2(FC) = 0.502); x-axis: chromosome position; y-axis: normalized snATAC-seq signal. **F)** Higher expression of SLC11A1 in nuclei from TS brains as compared to NC brains (adjusted p-value < 0.001, log2(FC) = 2.56). x-axis: expression level of SLC11A1; y-axis: density of number of nuclei.

We further profiled single-nuclei chromatin accessibility in 11 brains (sample from brain TS2 was excluded due to low number of nuclei passing QC) and annotated the 6 major cell types using the label transfer method from the snRNA-seq dataset (**Methods**). Out of 39,728 single nuclei which passed QC, 33% were classified as MSN, 2% as IN, 8% as AST, 47% as OL, 9% as MG, 1% as EC, and for 1% of cells their type was unidentified (**Figure 1F, G, H, Supplementary Table 4**,**5, Supplementary Figure 4**). Label transfer accuracy was 97.6%, as estimated comparing the multiome snRNA-seq and snATAC-seq data (**Supplementary Figure 5**). Together, the data indicate that using snRNA-seq and scATAC-seq we were able to map the principal cell types of the human striatum, including the relatively sparse populations of striatal interneurons. We observed no statistically significant differences in cell type composition between samples sequenced with the stand-alone techniques and samples sequenced with the multiome technique for both snRNA-seq and snATAC-seq data **(Supplementary Figure 6)**.

### Differential cell type composition and differentially expressed genes (DEGs) in patients with Tourette Syndrome

We used a paired t-test to compare proportions in each cell type between 5 TS and NC samples (one pair of samples (NC5/TS5) was excluded because of the low total count of nuclei passing QC in the paired control). We observed significantly fewer IN in the TS brains as compared to matched NC (2.4% vs 5.5% in TS vs NC, respectively, p-value < 0.001), with no significant difference in proportion for other cell types (**Figure 2A, Supplementary Figure 2, Supplementary Table 3**). This observation is concordant with previous immunocytochemistry results in a different cohort of postmortem brains (10-12).

To identify genes potentially involved in TS neurodevelopment, we assessed differentially expressed genes (DEGs) by comparing combined data from all single nuclei in TS and NC brains in each cell type (**Methods**). We observed between 300 and 600 DEGs in each cell type, with no statistically significant difference in the number of up and down-regulated DEGs **(Figure 2B, Supplementary Table 7)**. The signatures of up- and down-regulated genes were remarkably cell type-specific (**Supplementary Tables 8, 9**). In MSN, down-regulated DEGs were enriched in gene ontology (GO) annotation for mitochondrial function related pathways, such as mitochondrial inner membrane and ATP synthesis. In IN, down-regulated DEGs were enriched in plasma membrane/postsynaptic membrane annotations comprising GABA and AMPA/kainate glutamate post-synaptic receptors as well as several protocadherins, including PCDH17, PCDH19, PCDH7. There was no significant GO enrichment for down-regulated DEGs in other cell types. However, up-regulated DEGs were strongly enriched in immune response-related annotations in MG (**Figure 2C**). Downregulation in synaptic transmission-related and protocadherin-related gene modules and upregulation in immune response-related modules were also observed in a prior study of bulk RNA-seq expression performed in the basal ganglia of an independent cohort of TS subjects(12). Here we confirm these major signatures in single cells and assess their specific cellular origins – IN for synaptic transmission and protocadherin, MG for immune activation. To test the robustness of the DEGs identified in each cell type, we randomly down sampled each brain with 2000 nuclei **(Methods)**. As expected, we identified fewer number of DEGs in all cell types **(Supplementary Figure 7A, Supplementary Table 10);** IN in particular were strongly depleted in DEGs and displayed no significant GO enrichment, which is expected given the low proportion of IN in the striatum (3% of all cells, **Supplementary Fig. 2**). However, the same GO terms for down-regulated genes in MSN and up-regulated DEGs in MG remained significant **(Supplementary Figure 7B, Supplementary Table 11**,**12)**.

MG and MSN signatures represent different biological processes and consistently we see only 1 gene in common between the two signatures (69 DEGs in MSN signature and 33 DEGs in MG signature). To further understand whether the processes of mitochondrial disfunction in MSN and immunological activation of MG could be coupled, we correlated the two signatures across 5 pairs of brain samples (one pair was excluded due to low number of MGs). First, we ranked DEGs in MSN and MG in each pair and selected the 200 top ranked DEGs per cell type per pair. Then, we calculated enrichment scores for the selected DEGs in MSN in mitochondrial function-related GO terms and for the selected DEGs in MG in immune response-related GO terms (**Methods**). Signature scores were then correlated across brain pairs. This analysis revealed a statistically significant correlation between the two scores **(Figure 2D; Methods)**, suggesting potential cause-effect relationship between compromised oxidative metabolism in mitochondria of TS neurons and activation of the immune system in TS microglia.

### Differentially accessible regions in Tourette syndrome patients and co-accessibility networks

Across the 5 cell types we identified 339,857 peaks from snATAC-seq data, of which 89,234 were cell type specific **(Supplementary Table 13)**. About 68% of the identified snATAC-seq peaks overlapped with previously imputed bulk brain ATAC-seq peaks (27). Using the snATAC-seq data, we observed the same trend as in RNA-seq data of fewer IN in TS brains, however the trend didn’t reach statistical significance (**Figure 3A**). Among all peaks, 33% were residing within the exonic regions, 45% within intronic regions, 2% within the 2 kbp regions upstream and downstream of transcriptional start sites (TSS), which we named TSS-proximal peaks, and 20% within intergenic regions, which we named distal peaks (**Figure 3B**). We next performed GO annotation considering the genes in proximity to peaks. The GO biological process terms enriched for each cell type were consistent with the known function of the cell type; for example, synaptic transmission-related terms for MSN and IN, axon ensheathment and myelination for OL, and immune response signaling for MG (**Supplementary Figure 8**).

To investigate the differentially accessible regions, we compared the snATAC-seq peaks in TS brains vs NC in each major cell type (**Methods**). MSN and OL, which are the two most abundant cell types, exhibited the highest number of differentially accessible regions in TS vs NC (**Figure 3C, Supplementary Table 14, 15**).

Next, we investigated possible physical interactions between the identified open chromatin regions by analyzing their correlation across sub-groups of nuclei in each cell type (**Methods**). We focused on pairs of co-accessible peaks where one of the peaks was proximal to the TSS of a gene and the other one was a distal peak (**Methods**). Such pairs likely represented cis-regulatory interactions between putative enhancers and promoter regions and allowed us to investigate the correspondence between DEGs and putative enhancer activity in TS and NC (**Supplementary Table 16**). We identified 211 non-redundant DAP-DEG links, and computed a co-accessibility score, which is the correlation between a pair of peaks, with one overlapping with a DAP and another overlapping the promoter of a DEG. This was a minor fraction of identified links between distal peaks and DEG and between distal DAP and proximal peaks, likely reflecting the following: (i) sparseness of snATAC-seq data, i.e. higher propensity to capture promoters other than enhancers, (ii) limitation of the peak co-accessibility analysis relying on correlation between peaks using snATAC-seq data. For example, in MSN, we observed 3,396 distal peaks linked to DEGs, 846 distal DAPs linked to TSS-proximal peaks, and only 27 DAP-DEG links. Yet remarkably, we observed significant enrichments of the links between DAP-DEG links (i.e, those between a distal DAP and a DAP overlapping a DEG promoter) over total links in three cell types: MSN, OL and AST (**Supplementary Table 16**; p-value < 0.01 by Fisher’s exact test), revealing a relationship between gene expression and regulatory changes in TS. It is likely that no significance was observed IN and MG due to the few number of nuclei (<5%) analyzed.

To exemplify the significance of this observation, we investigated DAP-DEG links **(Supplementary Table 17)** and found that the promoter of the MG up-regulated gene solute carrier family 11 member 1 (SLC11A1) was co-accessible with a snATAC-seq distal peak **(Figure 3D,E)**. We observed that the snATAC-seq distal peak is more accessible in MGs of TS as compared to NC brains (**Figure 3E**), and that expression of SLC11A1 in TS is also higher (**Figure 3F**). This correlation suggests that the putative enhancer regulates SLC11A1 expression. The gene itself has multiple effects on macrophage activation and exerts an important role in immune response (28), which is concordant with the immune response-related gene up-regulation we observed in MG. In addition, we found that the MSN down-regulated gene Ubiquinol-Cytochrome C Reductase, Complex III Subunit XI (UQCR11) was co-accessible with less accessible distal snATAC-seq peak in MSNs (**Supplementary Fig. 9**). UQCR11 encodes for a binding factor of the ubiquinol-cytochrome c reductase complex, which forms part of the mitochondrial respiratory chain (29, 30). This is concordant with our observation of down-regulation of genes enriched in mitochondrial function related pathways from snRNA-seq in MSN.

## Discussion

In this study we perform a single cell-level transcriptome and epigenome analysis of the postmortem caudate nucleus in 6 pairs of adult TS and NC individuals and show that basal ganglia interneurons (IN) are decreased in number and display decreased expression in cell adhesion and synapse-related genes, while MSN show signs of metabolic stress and mitochondrial dysfunction in patients versus controls. At the same time, microglial cells are activated in TS, displaying increased expression of immune system-related genes. These results fit with a scenario where decreased tonic inhibition by IN upon MSN causes hyperactivity of the latter, along with signs of metabolic stress.

The picture is consistent with the established role of the basal ganglia in the storage of motor programs and modulation of motor behavior, where the globus pallidus (GPi) exerts tonic inhibition upon the thalamus and cortex, suppressing motor activity, and MSNs of the direct and indirect pathways integrate many inputs to modulate the GPi. The IN are a crucial component of such modulation as they receive input from the cortex and thalamus and exert prominent feed-forward inhibitory modulation upon the MSNs(26). Hence the decrease in IN number observed in TS is likely to be a crucial factor in the generation of tics.

Decreased number of IN (both cholinergic and GABAergic) has been previously reported in TS individuals (n=5 pairs) by immmunocytochemical studies (11). A subsequent bulk RNA-seq study performed in the striatum of 9 TS/NC pairs(12) suggested a decrease in inhibitory synaptic function (including a decrease in a number of synaptic channels and receptors) and increased immune activation but was unable to pinpoint the cellular origin of these alterations.

The current study on the single cell level, together with the previous evidence, provides a coherent picture of basal ganglia dysfunction by linking IN losses with potential hyper-excitability and bionergetic stress of MSN. Downregulation of membrane/synapses related genes in IN additionally suggest decreased arborization and connectivity, further impairing the function of these cells. The correlation between changes in gene expression and chromatin state suggests that some of these changes in transcriptome are driven by altered regulatory element activity. Further explorations of noncoding element activity in TS may provide insights into upstream causes of dysregulated gene expression and disfunction in patient’s brains which may have implications for treatment.

An important caveat of all postmortem studies is their descriptive nature, and particularly their inability to distinguish causes from consequences, which is noteworthy in cases of chronic brain dysfunction. One pertinent question is whether IN are dying or fail to be generated in TS. Our prior studies in iPSC-derived brain organoids from living TS and NC individuals suggested that BG IN are not generated in normal numbers due an aberrant response to the ventralizing morphogen Sonic hedgehog (SHH) in embryonic development (31), suggesting an early developmental origin for this IN loss, although we cannot exclude further losses later in life.

Implicating loss and dysfunction of BG interneurons in TS has implications for treatment. Current treatment strategies with antipsychotic drugs are not addressing the pathophysiology of the disorder. Our data suggest that compounds that increase interneuron neurotransmission may represent promising candidates.

## Acknowledgments

We acknowledge Livia Tomasini for technical advice on sample processing; the Tourette Association of America (TAA) for spearheading, organizing and supporting the postmortem brain collection for patients with Tourette syndrome. We thank the personnel at the Harvard Brain Tissue Resource Center/NIH NeuroBioBank for processing brain tissue; and Nancy Thompson for help with the clinical phenotyping. This work was supported in part by a fellowship grant from the Tourette Association of America (TAA) to L.F. and the National Institutes of Mental Health (R01 MH118453 to FMV). The Harvard Brain Tissue Resource Center is supported in part by the National Institutes of Health.

## Methods

### Subjects

TS brain samples of cases with refractory symptoms in adulthood, collected by the Tourette Association of America (TAA), were obtained from the Harvard Brain Tissue Resource Center. Samples of normal control subjects were obtained from the Harvard Brain Tissue Resource Center and from the Department of Pathology, Yale University (**Table S1**). Cases with known or suspected neurological diseases and prolonged hypoxia were excluded. There were no statistical differences in age, postmortem interval, or RNA quality between the TS and control groups (**Supplementary Table 1**).

### Sample preparation and nuclei isolation

Bulk brain tissue (caudate nucleus and putamen) was cut into 2mm cubes and was lysed by mechanical dissociation in 6ml lysis buffer (5.47g sucrose, 250ul CaCl_2_ (1M), 150ul Mg(Ac)_2_(1M), 10ul EDTA, 500ul Tris-HCL (1M), 17ul DTT (3M), 50ul Triton X in 50ml RNAse and DNAse free water). Nuclei were passed through a 40um diameter strainer and the lysate was transferred into a 15ml ultracentrifuge tube (Beckman). 9ml of ice cold sucrose solution (30.78g Sucrose, 150ul Mg(Ac)_2_ (1M), 500ul Tris-HCL (1M), 17ul DTT (3M) in 50ml RNAse and DNAse free water) was slowly added to the tube bottom. Nuclei were isolated by gradient ultracentrifugation under following conditions: 25,000rpm for 1h at 4° C.

### Standalone single nuclei RNA-seq and single nuclei ATAC-seq library preparation and sequencing

For single nuclei RNA-seq: After ultracentrifugation, supernatant was aspirated and the nuclei resuspended in 1x PBS and RNAse inhibitor (Roche, RNAse inhibitor concentration according to suppliers’ recommendation). For single nuclei ATAC-seq: After ultracentrifugation, supernatant was aspirated and the nuclei (3,080-7,700 nuclei/ul as recommended by 10X Genomics) were resuspended in 1X nuclei buffer (10X genomics). Samples were submitted to the Yale Center for Genome Analysis core for library preparation (10,000 nuclei/sample) using Chromium Single Cell ATAC reagent kit (10X Genomics).

### Multiome single nuclei RNA-seq and ATAC-seq library preparation and sequencing

After ultracentrifugation, the supernatant was aspirated and the nuclei were resuspended in ice cold 1X nuclei buffer (10X nuclei buffer stock, provided by 10X genomics) and nuclei were handled according to supplier’s recommendations. Samples were passed through a 40um diameter strainer. Nuclei were counted (Countess II, Thermo Fisher) and samples were submitted to Yale Center for Genome Analysis core for library preparation (10,000 nuclei/sample) using the Chromium Next GEM Single Cell Multiome ATAC + Gene Expression kit (10X Genomics) .

### snRNA-seq data processing, clustering and annotation

The gene count matrices for standalone and multiome snRNA-seq data were calculated using cellranger-6.0.1 with gex-GRCh38 as reference genome and genome annotation. The ribosomal genes were then removed from the count matrices. The single nuclei libraries with an extremely low number of genes (<200) and an extremely high number of genes (>mean+2*standard deviation) were removed from the analysis. All 12 samples were then merged using Seurat 5.0.1 (32-34) merge function. Potential doublets in the snRNA-seq data were removed using solo (35). The single nuclei libraries with too many mitochondrial gene expressions (>5%) were removed from the analysis. Expression of all genes for all the single nuclei were then normalized and integrated using the top 2000 expressed genes with Seurat normalization and integration functions. Dimension reduction was performed using UMAP with 50 dimensions as input features. Each cluster was annotated using literature (25, 36) and public datasets (https://www.brainspan.org) and clusters were combined into major cell types.

### snATAC-seq data processing, clustering and label transfer

The fragment count matrices for standalone and multiome snATAC-seq data were calculated using cellranger ARC-2.0.0 with arc-GRCh38 as reference genome and genome annotation. The single nuclei libraries with an extremely low number of fragments (<3000) or an extremely high number of fragments (>20000) were removed. The low quality single nuclei with <15% fragments in the predicted peak regions and single nuclei with nucleosome signal < 2 or > 4 were removed based on the suggestion provided by Signac (37). The count matrices of 12 samples were merged and integrated using features with more than 10 total counts. The integrated count matrix was then normalized and integrated for dimension reduction using UMAP with 30 dimensions as the input features. The common anchors (genes) were then identified between the snRNA-seq and snATAC-seq count matrices for label transferring using Signac(37).

### snRNA-seq differential gene expression

Genes present in at least 10% of cells in either TS patients or controls were considered in the differential gene expression analysis. Differential gene expression analysis was performed using the glmgampoi (38) R package v1.13.2. For each cell type, pairs with one sample having an extremely low number of single nuclei were not considered in the analysis (pairs included in analyses in different cell types: MSN: all pairs; IN: pair 1,2,3,19; AST: all pairs; OLI: all pairs; MG: pair 2,3,4,19). Genes were considered as differentially expressed in a specific cell type when exhibiting an adjusted p-value < 0.01 and an absolute log2 fold change > 0.5. GO term analysis was performed using DAVID Functional Annotation (39). To check for robustness, 2,000 single nuclei were randomly sampled from all 12 brain samples. If there are fewer than 2,000 single nuclei in one sample, all single nuclei in that sample were considered. With the randomly sampled nuclei, the same differentially gene expression analysis was performed using glmgampoi (38).

### Correlation analysis between MSN DEGs enriched signature and MG DEGs enriched signature

Five ranked lists of DEGs were identified for MSN and MG between pair 2,3,4,5, 19 (pair 1 was excluded due to low number of MGs) using the glmgampoi (38) R package v1.13.2. The enrichment score for the DEGs within a signature in the top 200 DEGs in each pair was calculated using the GSEA python package (40). The R^2 and p-value for the correlation analysis for 5 pairs of enrichment scores were calculated using Pearson correlation test.

### snATAC-seq differentially accessible peaks and co-accessibility analysis

Peaks were called using CallPeaks function in Signac v1.12.0 (37) with MACS2 (41). Differentially active peaks were then identified using the FindMarkers function by comparing each cell type to the rest of the cells as well as by comparing the single nuclei in TS patients to the single nuclei in controls. Peaks were considered as differentially accessible when their adjusted p-value was <= 0.1 and the log2 fold change was > 0.2. GO analysis was performed using GREAT (42, 43). Co-accessible sites were identified using Cicero v1.3.9 (44) with pairs ranking at the top 5% co-accessibility score for considering significant links between two peaks. The links were then overlapped with transcriptional start sites (TSS) of genes to identify TSS-proximal peaks and potential gene-enhancer links.

