## Supplementary Figures for "Interneuron loss and microglia activation by transcriptome analyses in the basal ganglia of Tourette syndrome"

### Supplemental Information

Yifan Wang[1], Liana Fasching[2], Feinan Wu[2], Anita Huttner[3], Sabina Berretta[4], Rosalinda Roberts[5], James F. Leckman[2], Alexej Abyzov[1], Flora M. Vaccarino[2,6,7]

[1]Department of Quantitative Health Sciences, Center for Individualized Medicine, Mayo Clinic, Rochester, MN 55905, USA
[2]Child Study Center, Yale University, New Haven, CT 06520, USA
[3]Department of Pathology, Yale University, New Haven, CT 06520, USA
[4]McLean Hospital, Harvard Medical School, Belmont, MA, 02478, USA
[5]Department of Psychiatry and Behavioral Neurobiology, University of Alabama at Birmingham, USA
[6]Department of Neuroscience, Yale University, New Haven, CT 06520, USA
[7]Yale Kavli Institute for Neuroscience, New Haven, CT 06520, USA

This supplement contains:

Supplementary Figures 1 through 8

Supplementary Tables 1 through 6 and 16

Additionally, the following Supplementary tables are separately uploaded as excel files:

Supplementary Table 7 (Stable 7). Differentially expressed genes (DEG) between TS and NC brains in each cell type.
Supplementary Table 8 (Stable 8). GO terms enriched in down regulated DEGs.
Supplementary Table 9 (Stable 9). GO terms enriched in up regulated DEGs.
Supplementary Table 10 (Stable 10). Differentially expressed genes in each cell type with random sampled single nuclei.
Supplementary Table 11 (Stable 11). GO terms enriched in down regulated DEGs with down sampled nuclei.
Supplementary Table 12 (Stable 12). GO terms enriched in up regulated DEGs with down sampled nuclei.
Supplementary Table 13 (Stable 13). snATAC-seq peaks identified from 12 brains.
Supplementary Table 14 (Stable 14). Less accessible peaks in TS brains.
Supplementary Table 15 (Stable 15). More accessible peaks in TS brains.
Supplementary Table 17 (Stable 17). DAP-DEG co-accessible peak pairs in each cell type.

A

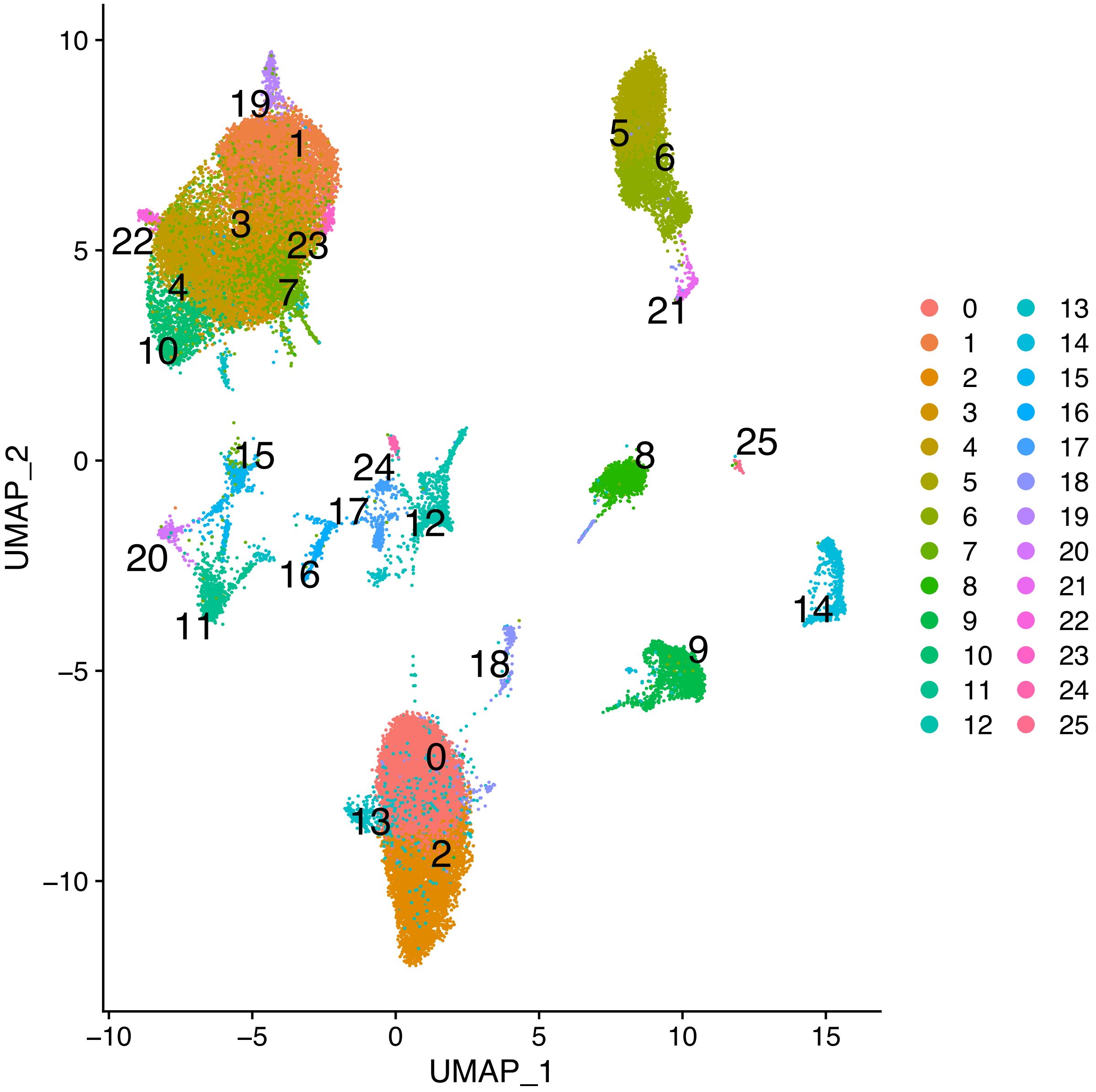

UMAP_2
UMAP_1
10
5
0
-5
-10
-10
-5
0
5
10
15
19
1
3
22
23
4
7
10
5
6
21
15
24
17
12
20
16
11
8
25
14
9
18
0
13
2
0 1 2 3 4 5 6 7 8 9 10 11 12 13 14 15 16 17 18 19 20 21 22 23 24 25

B

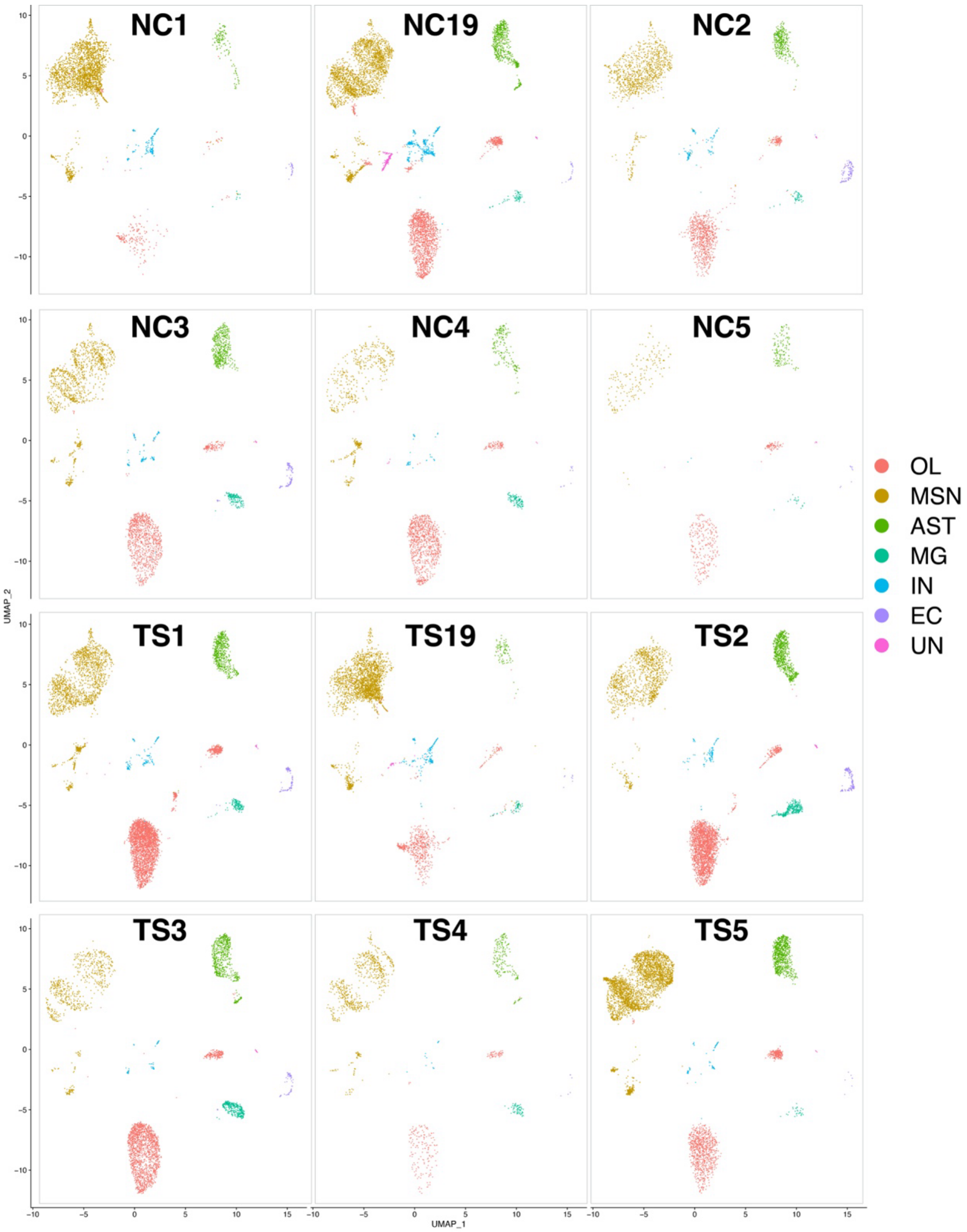


**Supplementary Figure 1.** UMAP clustering of nuclei from 12 brains based on snRNA-seq data. **A)** 26 clusters for the entire set of 12 brains. **B)** Clusters were grouped into 6 major cell types and their distribution is shown across the 12 brains. Color: different cell types.

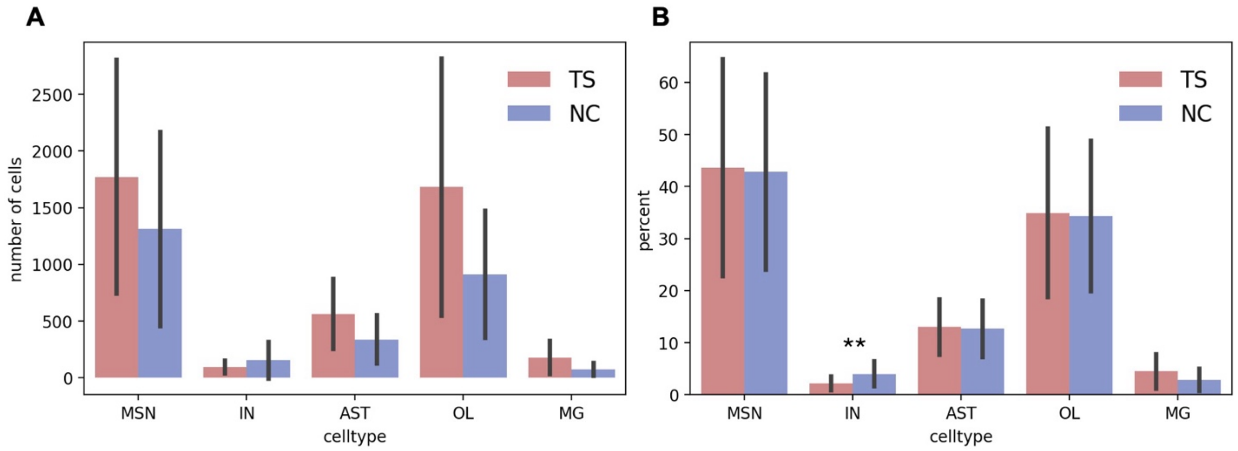


**Supplementary Figure 2. Cell type composition of TS and NC brains in snRNA-seq. A)** Number of cells in each cell type in 6 pairs of samples. **B)** percentage with respect to library size of each cell type in 6 pairs of samples. Red: TS patients; blue: controls. Error bar: standard deviation; **: p-value < 0.001 by paired t-test.

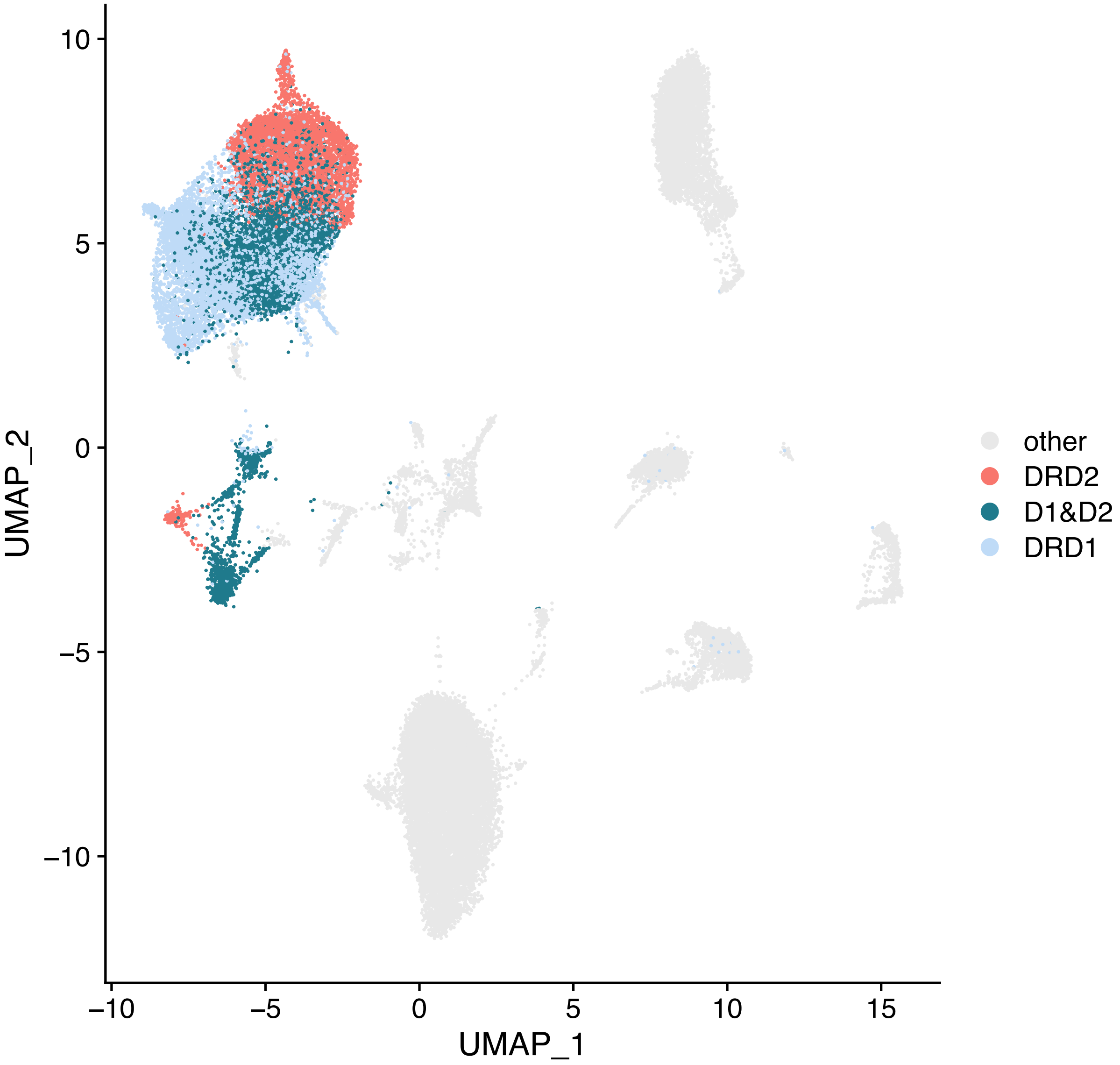


**Supplementary Figure 3. Cell type composition of the MSN cluster in the entire dataset (12 brains).** DRD1: dopamine receptor 1 (DRD1)-MSN1; DRD2: dopamine receptor 2 (DRD2)-MSN2; D1&D2: mixture of DRD1-MSN1 and DRD2-MSN2; other: other cell types.

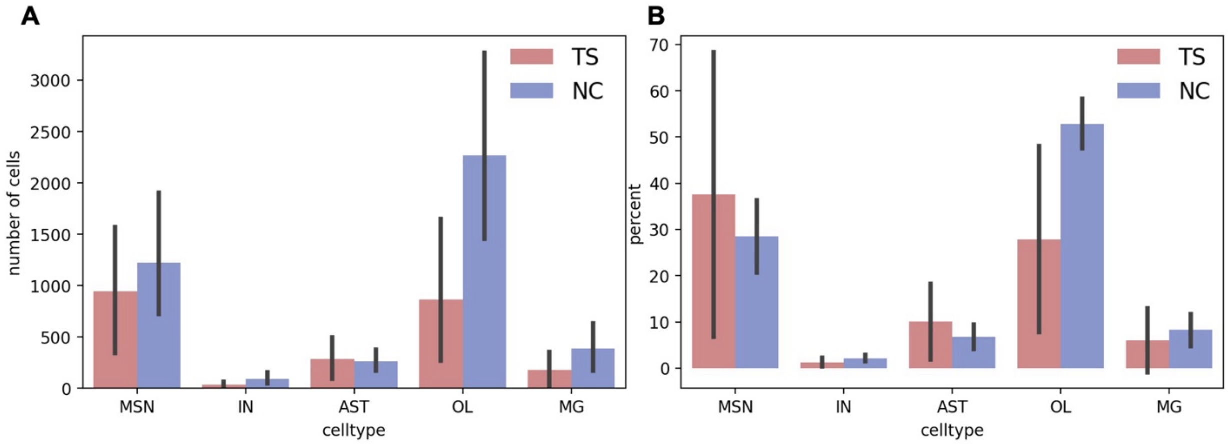


**Supplementary Figure 4. Cell type composition of TS and control brains in snATAC-seq. A)** Number of cells in each cell type in 6 pairs of samples. **B)** percentage of each cell type in 6 pairs of samples. Red: TS patients; blue: controls. Error bar: standard deviation.

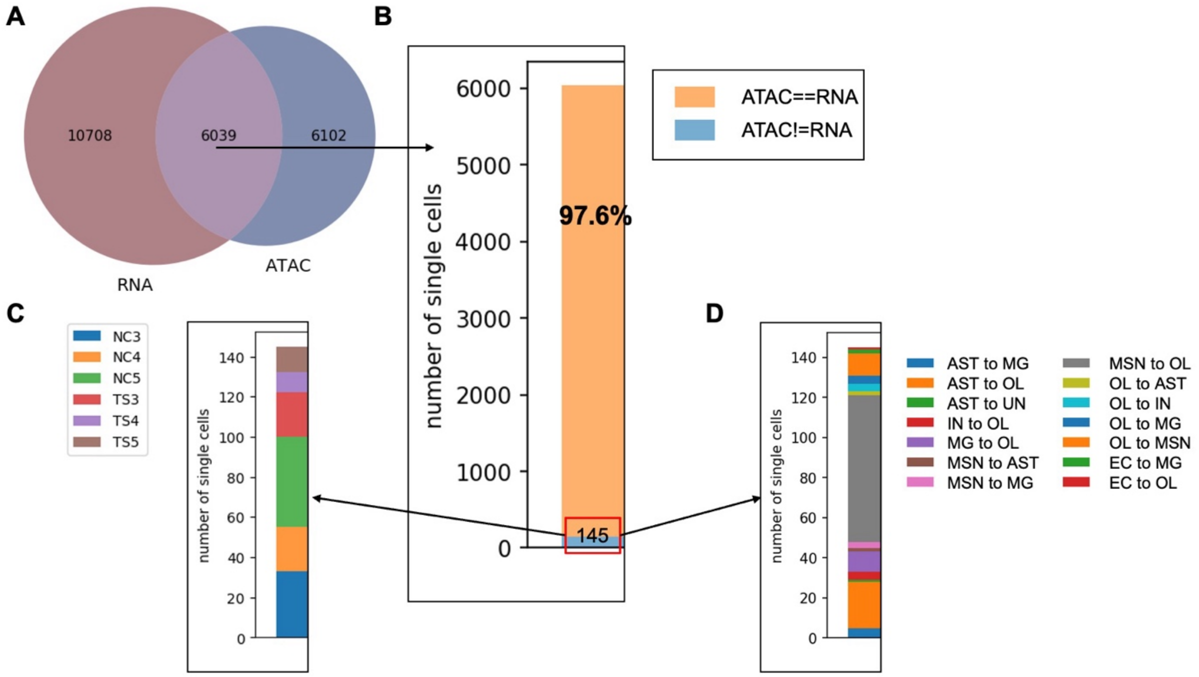


**Supplementary Figure 5. Comparison of cell type predicted using label transfer and those predicted with multiome barcodes. A)** Number of nuclei in snRNA-seq (dark red), snATAC-seq (dark blue), and the nuclei with the same barcode from multiome data (Light purple). **B)** Number of single nuclei with the same (orange) or different (blue) cell types from label transfer prediction and multiome. **C)** Sample distribution of the 2.4% nuclei with different cell type predictions. **D)** Cell type mislabeling distribution of the 2.4% nuclei.

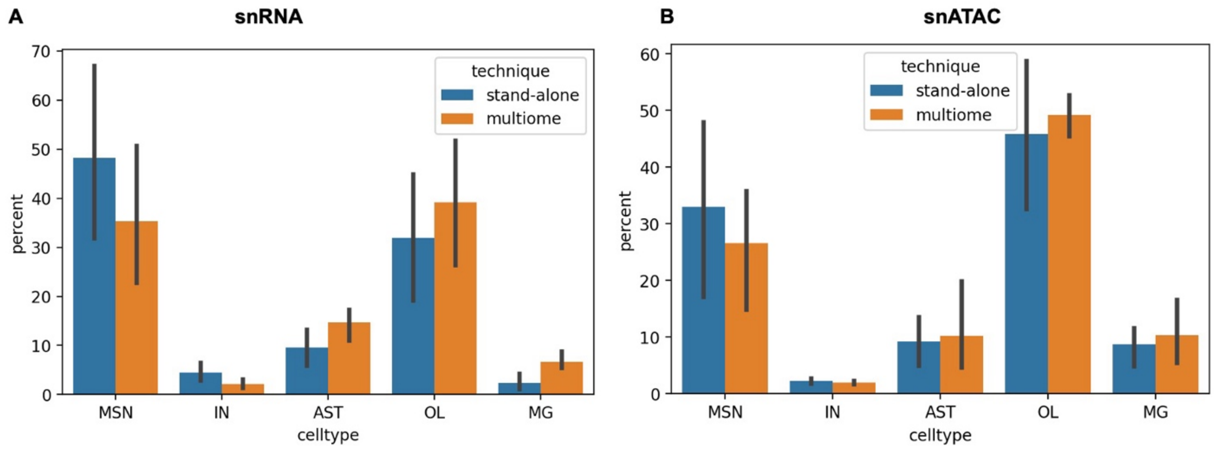


**Supplementary Figure 6. Cell type composition comparison between stand-alone and multiome techniques in snRNA-seq and snATAC-seq. A)** Percentage of each cell type for the 12 brains in stand-alone snRNA-seq and multiome snRNA-seq. **B)** Percentage of each cell type for 12 brains in stand-alone snATAC-seq and multiome snATAC-seq. Error bar: standard deviation.

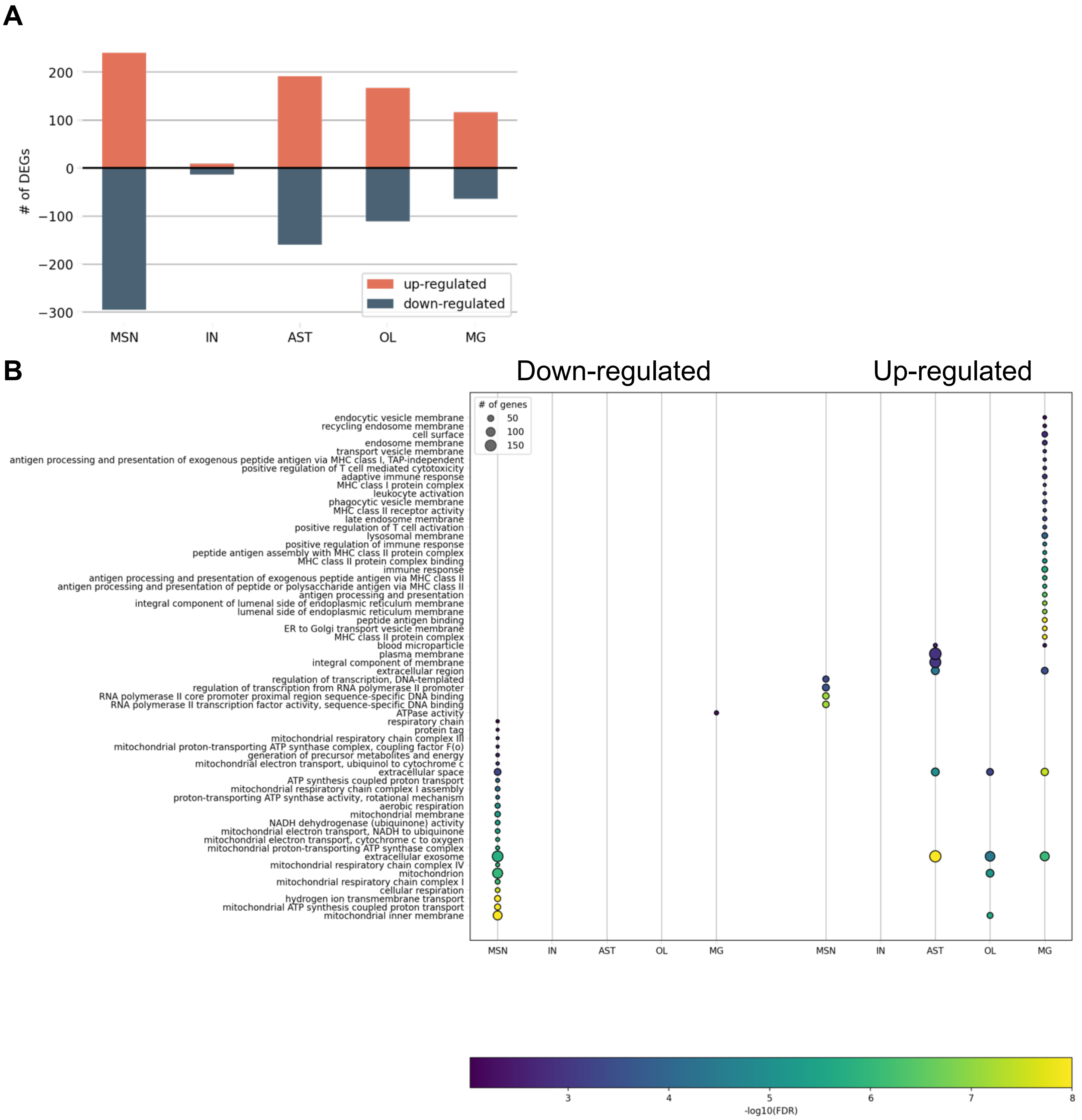


**Supplementary Figure 7.** DEGs after randomly down sampling each brain to 2000 nuclei. **A)** Number of differentially expressed genes in each cell types comparing TS to NC brains. **B)** GO terms enrichment for DEGs comparing TS and NC after random down sampling. y-axis: enriched GO terms; x-axis: cell types. The size of a circle indicates the number of DEGs in the enriched GO term, while the color of a circle corresponds to -log10(FDR) of the enriched GO term. GO terms were ordered based on the p-value in each cell type.

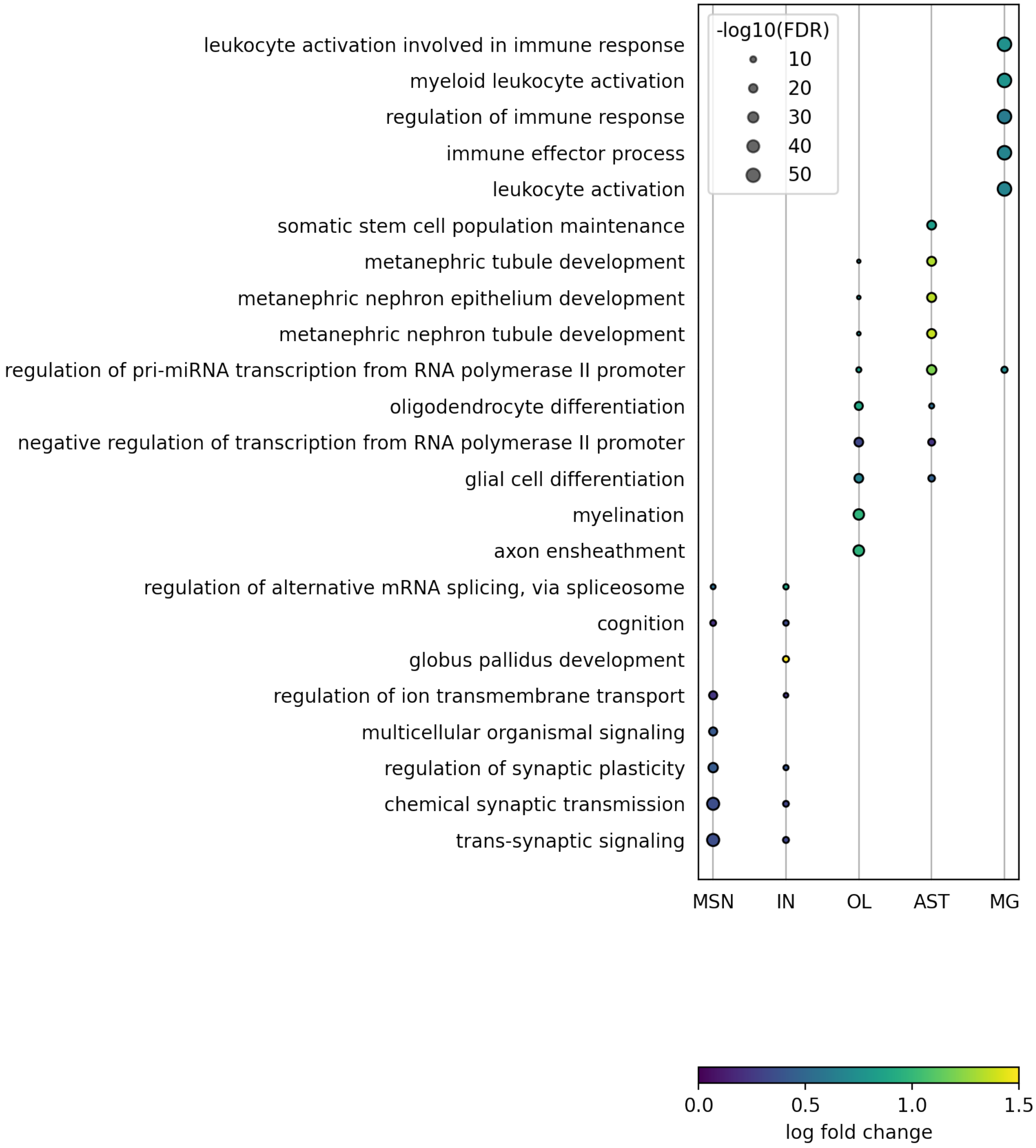


**Supplementary Figure 8.** GO terms enrichment for differentially accessible peaks in different cell types. y-axis: enriched GO terms; x-axis: cell types. The color of a circle indicates the log2 fold change of the enriched GO term, while the size of a circle corresponds to -log10(FDR) of the enriched GO term. GO terms were ordered based on the p-value in each cell type.

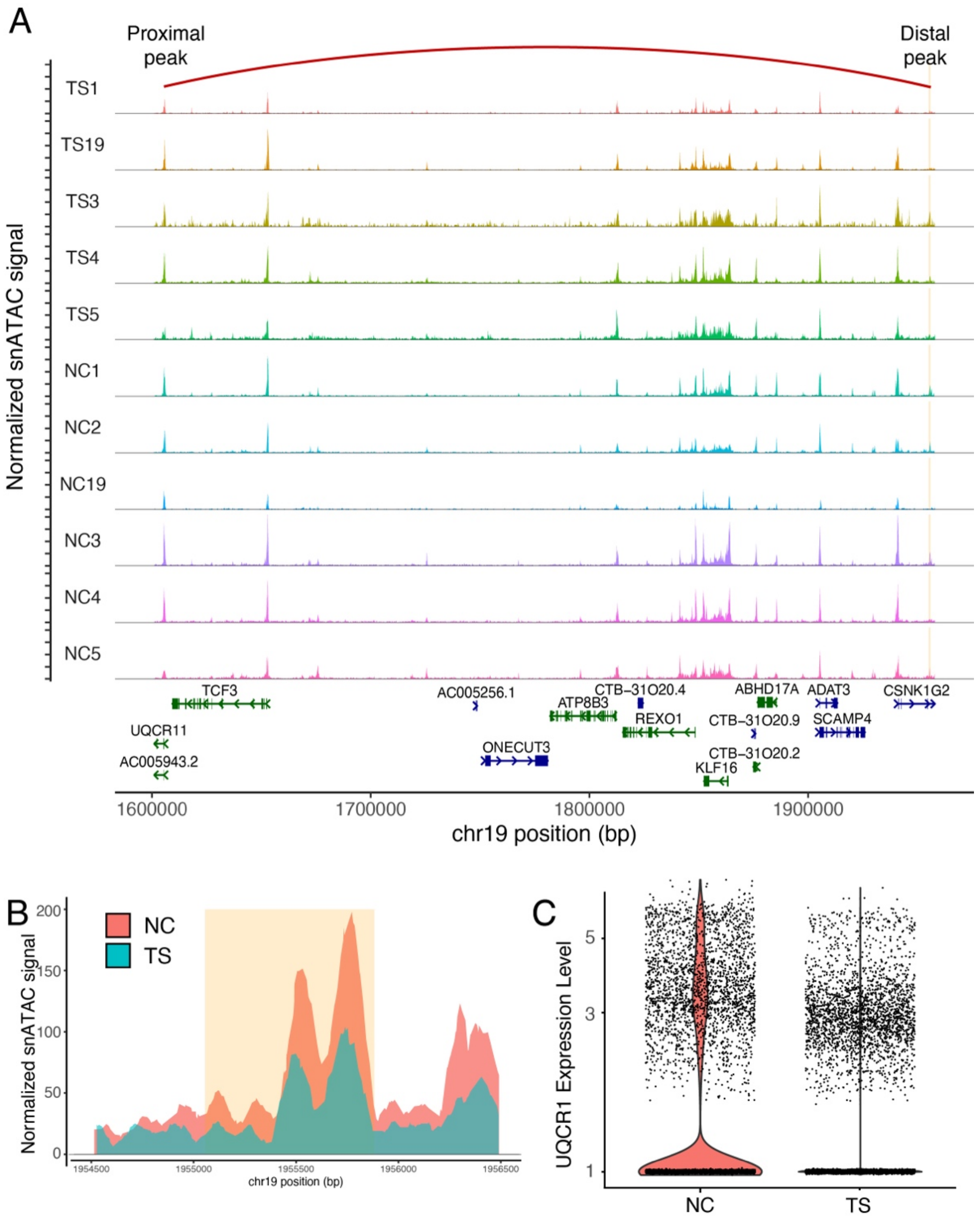


**Supplementary Figure 9.** DAP-DEG pair between the potential regulatory region and the promoter of UQCR11 in MSN. **A)** Down-regulation of UQCR11 associated with less accessible peaks in TS brains. x-axis: genome coordinates; y-axis: aggregated normalized snATAC-seq signal in MSNs of each brain; orange areas: snATAC-seq distal peak co-accessible with UQCR11 promoter; red arches: predicted co-accessible peak pair. **B)** Less accessible distal snATAC-seq DAP in TS brains compared to NC brains co-accessible with UQCR11 promoter; x-axis: chromosome coordinates; y-axis: normalized snATAC-seq signal in MSNs. **C)** UQCR11 down-regulation in TS brains; x-axis: NC or TS brains; y-axis: UQCR11 expression level in each single nucleus of MG.

**Supplementary Table 1: human brain sample description and metadata**

| SPECIMEN | group | sample name sequencing library | Experiment | age | sex | PMI |
|---|---|---|---|---|---|---|
| 5627 | Tourette | TS1 | separate sn-RNA seq and sn-ATAC seq | **42** | **m** | **25h** |
| 4587 | Tourette | TS2 | separate sn-RNA seq and sn-ATAC seq | **52** | **f** | **5h** |
| 7099 | Tourette | TS19 | separate sn-RNA seq and sn-ATAC seq | **64** | **m** | **20h** |
| 13082 | Control | NC1 | separate sn-RNA seq and sn-ATACs eq | **41** | **m** | **27.4h** |
| 7087 | Control | NC2 | separate sn-RNA seq and sn-ATAC seq | **52** | **f** | **21.51h** |
| FV05 | Control | NC19 | separate sn-RNAs eq and sn-ATAC seq | **66** | **m** | **36h** |

| SPECIMEN | group | sample name sequencing library (sn-RNA seq/sn-ATAC seq) | Experiment | age | sex | PMI |
|---|---|---|---|---|---|---|
| AN11896/B7381 | Tourette | TS3_CDPT_3P/TS3_CDPT_ATAC | Multiome | **35** | **m** | **30h** |
| AN18879/B4790 | Tourette | TS4_CDPT_3P/TS4_CDPT_ATAC | Multiome | **35** | **m** | **30h** |
| AN05094/B4184 | Tourette | TS5_CDPT_3P/TS5_CDPT_ATAC | Multiome | **77** | **m** | **11h** |
| 13138 | Control | NC3_CDPT_3P/NC3_CDPT_ATAC | Multiome | **40** | **m** | **31h** |
| 5715 | Control | NC4_CDPT_3P/NC4_CDPT_ATAC | Multiome | **35** | **m** | **27h** |
| FV03 | Control | NC5_CDPT_3P/NC5_CDPT_ATAC | Multiome | **66** | **m** | **39** |

**Supplementary Table 2. Number of cells in each cell type for every sample in snRNA-seq**

| | MSN | IN | OL | AST | MG | EC | UN |
|---|---|---|---|---|---|---|---|
| NC3 | 1188 | 122 | 1164 | 518 | 171 | 125 | 11 |
| TS3 | 716 | 54 | 1973 | 684 | 381 | 69 | 22 |
| NC4 | 564 | 47 | 1030 | 179 | 118 | 11 | 6 |
| TS4 | 696 | 14 | 219 | 198 | 68 | 6 | 0 |
| NC5 | 178 | 2 | 320 | 149 | 18 | 10 | 5 |
| TS5 | 3553 | 75 | 1143 | 898 | 32 | 7 | 3 |
| NC1 | 2259 | 128 | 192 | 118 | 8 | 23 | 4 |
| TS1 | 1907 | 149 | 3508 | 628 | 148 | 110 | 27 |
| NC2 | 1096 | 122 | 893 | 357 | 50 | 132 | 7 |
| TS2 | 1210 | 95 | 2588 | 864 | 375 | 193 | 53 |
| NC19 | 2578 | 504 | 1876 | 708 | 68 | 25 | 379 |
| TS19 | 2553 | 180 | 657 | 93 | 59 | 8 | 34 |

**Supplementary Table 3. Percentage of cells in each cell type for every sample in snRNA-seq**

| | MSN | IN | OL | AST | MG | EC | UN |
|---|---|---|---|---|---|---|---|
| NC3 | 36.01 | 3.70 | 35.28 | 15.70 | 5.18 | 3.79 | 0.33 |
| TS3 | 18.36 | 1.38 | 50.60 | 17.54 | 9.77 | 1.77 | 0.56 |
| NC4 | 28.85 | 2.40 | 52.69 | 9.16 | 6.04 | 0.56 | 0.31 |
| TS4 | 57.95 | 1.17 | 18.23 | 16.49 | 5.66 | 0.50 | 0.00 |
| NC5 | 26.10 | 0.29 | 46.92 | 21.85 | 2.64 | 1.47 | 0.73 |
| TS5 | 62.21 | 1.31 | 20.01 | 15.72 | 0.56 | 0.12 | 0.05 |
| NC1 | 82.69 | 4.69 | 7.03 | 4.32 | 0.29 | 0.84 | 0.15 |
| TS1 | 29.44 | 2.30 | 54.16 | 9.70 | 2.29 | 1.70 | 0.42 |
| NC2 | 41.25 | 4.59 | 33.61 | 13.44 | 1.88 | 4.97 | 0.26 |
| TS2 | 22.50 | 1.77 | 48.12 | 16.07 | 6.97 | 3.59 | 0.99 |
| NC19 | 42.00 | 8.21 | 30.56 | 11.53 | 1.11 | 0.41 | 6.17 |
| TS19 | 71.23 | 5.02 | 18.33 | 2.59 | 1.65 | 0.22 | 0.95 |

**Supplementary Table 4. Number of cells in each cell type for every sample in snATAC-seq**

| | MSN | IN | OL | AST | MG | EC | UN |
|---|---|---|---|---|---|---|---|
| NC3 | 918 | 68 | 1437 | 114 | 310 | 80 | 27 |
| TS3 | 122 | 36 | 733 | 413 | 334 | 23 | 18 |
| NC4 | 542 | 27 | 961 | 129 | 119 | 0 | 2 |
| TS4 | 650 | 32 | 878 | 94 | 74 | 1 | 1 |
| NC5 | 997 | 73 | 1336 | 298 | 52 | 2 | 5 |
| TS5 | 1155 | 0 | 32 | 48 | 0 | 0 | 0 |
| NC1 | 2809 | 233 | 4201 | 245 | 918 | 51 | 34 |
| TS1 | 1551 | 63 | 801 | 464 | 86 | 2 | 3 |
| NC2 | 1585 | 50 | 2954 | 488 | 400 | 65 | 44 |
| TS2 | 0 | 0 | 0 | 0 | 0 | 0 | 0 |
| NC19 | 497 | 115 | 2698 | 305 | 525 | 2 | 85 |
| TS19 | 2200 | 95 | 2734 | 701 | 564 | 3 | 16 |

**Supplementary Table 5. Percentage of cells in each cell type for every sample in snATAC-seq**

| | MSN | IN | OL | AST | MG | EC | UN |
|---|---|---|---|---|---|---|---|
| NC3 | 31.08 | 2.30 | 48.65 | 3.86 | 10.49 | 2.71 | 0.91 |
| TS3 | 7.27 | 2.14 | 43.66 | 24.60 | 19.89 | 1.37 | 1.07 |
| NC4 | 30.45 | 1.52 | 53.99 | 7.25 | 6.69 | 0.00 | 0.11 |
| TS4 | 37.57 | 1.85 | 50.75 | 5.43 | 4.28 | 0.06 | 0.06 |
| NC5 | 36.08 | 2.64 | 48.35 | 10.79 | 1.88 | 0.07 | 0.18 |
| TS5 | 93.52 | 0.00 | 2.59 | 3.89 | 0.00 | 0.00 | 0.00 |
| NC1 | 33.08 | 2.74 | 49.48 | 2.89 | 10.81 | 0.60 | 0.40 |
| TS1 | 52.22 | 2.12 | 26.97 | 15.62 | 2.90 | 0.07 | 0.10 |
| NC2 | 28.37 | 0.90 | 52.88 | 8.74 | 7.16 | 1.16 | 0.79 |
| TS2 | 0.00 | 0.00 | 0.00 | 0.00 | 0.00 | 0.00 | 0.00 |
| NC19 | 11.76 | 2.72 | 63.83 | 7.22 | 12.42 | 0.05 | 2.01 |
| TS19 | 34.85 | 1.50 | 43.31 | 11.10 | 8.93 | 0.05 | 0.25 |

**Supplementary Table 6. Markers for cell type annotation for BG cell types and cluster feature plot**

| gene | subclass celltype | Gene full name |
|---|---|---|
| **ChAT** | cholinergic IN | Choline O-Acetyltransferase |
| **SLC5A7** | cholinergic IN | Solute Carrier Family 5 Member A7 |
| **SLC18A3** | cholinergic IN | Solute Carrier Family 18 Member A3 |
| **CHRM2** | cholinergic IN | Cholinergic Receptor Muscarinic 2 |
| **GBX2** | cholinergic IN | Gastrulation Brain Homeobox 2 |
| **GAD1** | GABA IN | Glutamate Decarboxylase 1 |
| **DLX1** | GABA IN | Distal-Less Homeobox 1 |
| **DLX2** | GABA IN | Distal-Less Homeobox 2 |
| **DLX5** | GABA IN | Distal-Less Homeobox 5 |
| **DLX6** | GABA IN | Distal-Less Homeobox 6 |
| **LHX6** | MGE-GABA IN | LIM Homeobox 6 |
| **NOS1** | MGE-GABA IN | Nitric Oxide Synthase 1 |
| **NPY** | POA- GABA IN | Neuropeptide Y |
| **SST** | MGE- GABA IN | Somatostatin |
| **CALB2** | CGE-GABA IN | Calbindin 2/Calretinin |
| **PVALB** | MGE- GABA IN | Parvalbumin |
| **KCNC1** | MGE-GABA IN | Potassium Voltage-Gated Channel Subfamily C Member 1 |
| **CALB1** | POA-IN | Calbindin |
| **VIP** | CGE-IN | Vasoactive Intestinal Peptide |
| **RELN** | CGE-IN | Reelin |
| **FOXP1** | LGE-MSN | Forkhead Box Protein P1 |
| **PPP1R1B** | LGE-MSN | DARPP-32/ Protein Phosphatase 1 Regulatory Inhibitor Subunit 1B |
| **DRD1** | LGE-MSN1 | Dopamine Receptor 1 |
| **DRD2** | LGE-MSN2 | Dopamine Receptor 2 |
| **PENK** | LGE-MSN2 | Proenkephalin |
| **PDYN** | LGE-MSN1 | Prodynorphin |
| **TAC1** | LGE-MSN1 | Tachykinin Precursor 1/Substance P |
| **OLIG1** | OL | Oligodendrocyte Transcription Factor 1 |
| **OLIG2** | OL | Oligodendrocyte Transcription Factor 2 |
| **GFAP** | AST | Glial fibrillary acidic protein |
| **AQP4** | AST | Aquaporin-4 |
| **CD74** | MG | A major histocompatibility complex class II (MHCII) interacting molecule expressed in microglia |

| | | |
|---|---|---|
| **RGS10** | MG | Regulator Of G Protein Signaling 10 |

Annotation of basal ganglia interneurons (IN) and Medium Spiny Neurons (MSN) according to their gene expression and embryonic origin in the ganglionic eminences (GE) of the basal forebrain (1, 2). Cholinergic neurons have mixed origin from the MGE and the anterior preoptic area (POA) of the hypothalamus.

IN=interneuron; MGE= medial ganglionic eminence; LGE= lateral ganglionic eminence CGE= caudal ganglionic eminence; POA= anterior preoptic area

**Supplementary Table 16. Number of co-accessible peak pairs between potential regulatory regions (distal peaks) and promoters (TSS-proximal peaks).**

| | MSN | IN | AST | OL | MG |
|---|---|---|---|---|---|
| Total # of links | 239,628 | 133,027 | 194,648 | 438,485 | 114,404 |
| # of links between a DEG and a distal peak | 3,396 | 1,588 | 3,472 | 4,216 | 1,134 |
| # of links between a DAP and a proximal peak | 846 | 3 | 547 | 3897 | 549 |
| DEG-DAP links | 27 | 0 | 22 | 151 | 11 |
| Fisher Exact test p-value | 0.0001 | NA | 0.0005 | 1.4e-45 | 0.03 |

An example for Fisher Exact test for MSN:

| | DEG | Not DEG |
|---|---|---|
| DAP | 27 | 846–27 = 819 |
| Not DAP | 3,396–27 = 3,369 | 239,628-3,369-819-27 =235,413 |


1. Lim L, Mi D, Llorca A, Marin O (2018): Development and Functional Diversification of Cortical Interneurons. *Neuron*. 100:294-313.
2. Crittenden JR, Graybiel AM (2011): Basal Ganglia disorders associated with imbalances in the striatal striosome and matrix compartments. *Front Neuroanat*. 5:59.
